# A tipping point in West Antarctica in 2023’s circumpolar 5-sigma event? a glimpse from the emperor penguin’s perspective

**DOI:** 10.1101/2023.09.28.559910

**Authors:** Lucas Krüger, Juliana A. Vianna, Céline Le Bohec, César A. Cárdenas

**Affiliations:** Departamento Científico, Instituto Antártico Chileno (INACH), Punta Arenas, Plaza Muñoz Gamero 1055, Chile; Millennium Institute of Biodiversity of Antarctic and Subantarctic Ecosystems (BASE), Santiago, Chile; Pontificia Universidad Católica de Chile, Facultad de Ciencias Biológicas, Departamento de Ecología, Instituto para el Desarrollo Sustentable, Av. Bernardo O’Higgins 340, Santiago, Chile; Millennium Institute Center for Genome Regulation (CRG), Santiago, Chile; Millennium Nucleus of Patagonian Limit of Life (LiLi), Santiago, Chile; Université de Strasbourg, CNRS, IPHC UMR 7178, Strasbourg, France; CEFE, Université de Montpellier, CNRS, EPHE, IRD, Montpellier, France; Centre Scientifique de Monaco, Département de Biologie Polaire, Monaco, Principality of Monaco

**Keywords:** Antarctic - Amundsen Sea, *Aptenodytes forsteri*, Belingshausen Sea, Seabirds, Sea Ice

## Abstract

Emperor penguins are one of the seabird species most vulnerable to climate change in Antarctica, as they depend on stable sea ice for breeding and molting. Recent extremes of warming have taken to the lowest records of sea ice in West Antarctica in 2022 and 2023, raising concerns to the extent that emperor penguins have been affected. We evaluated historical satellite data (1985-2022) of sea ice around the 10 emperor penguin breeding colonies, identified so far along the coast of the Amundsen and Bellingshausen Seas, and compared them to recent sea ice cover (2022-2023) to identify where and when critical thresholds of sea ice cover have been crossed. We found that the sea ice retreated below the critical thresholds in 2022-2023 during the chick-rearing and in some cases failed to return at the start of the following breeding season, hence likely affecting 9 of the 10 emperor penguin colonies in the region. This is an unprecedented climatic phenomena affecting breeding habitat of emperor penguin colonies over a large area, potentially affecting numerically important colonies in West Antarctica.

## Introduction

Antarctic sea ice has been following a downward trend that started about one decade ago (Eayrs et al. 2021) due to warming and the increased frequency of intense heat waves or warm atmospheric rivers (Turner et al. 2021; Hepworth et al. 2022; Wille et al. 2022). As a consequence, Antarctic sea ice cover in 2022 and even more so in 2023, with the below 5-sigma threshold, has been at its lowest since the start of measurements, particularly in West Antarctica (Yadav et al. 2022; Liu et al. 2023; Scott and Scambos 2023).

Emperor penguins have been identified as one of the most vulnerable species under the projected climate change scenarios in Antarctica (Lee et al. 2022). Critical months for the breeding season of emperor penguins include August to mid-December (Handley et al. 2021). Population dynamics of this species are heavily linked to sea ice (Croxall 2002; Jenouvrier et al. 2014), as they are dependent on stable sea (fast) ice or ice shelves for breeding and molting (Kooyman et al. 2000, 2004; Fretwell et al. 2014; Zitterbart et al. 2014). Collapse of sea ice during late crèche stage (October to December) have resulted in high mortality in some colonies, with posterior slow recovery (Croxall 2002; Trathan et al. 2011). As a consequence, breeding sites seem to have been abandoned after consecutive years of unsuitable conditions (Trathan et al. 2011; Fretwell and Trathan 2019).

The recent records of low sea ice cover in west Antarctica during summer (Liu et al. 2023) poses a serious concern for emperor penguins (Fretwell et al. 2023). Rapid increase in loss rate of stable sea ice could represent a substantial decrease in the availability of suitable breeding habitats (Trathan et al. 2020). Emperor penguin colonies in the Bellingshausen and Amundsen seas have been forecasted to experience steep decreases within a period of a few decades (Jenouvrier et al. 2021), and it is possible that 2022 and 2023 events mark a quickening of that process (Fretwell et al. 2023). With the recent reports of intense warming taking to the lower levels of sea ice ever recorded in the Amundsen and Bellingshausen Sea regions, we evaluated here the condition of sea ice at emperor penguin colony sites by using a 39-years interval to calculate local sea ice daily means and identifying periods between January 2022 and June 2023 when sea ice was below the historical 20th percentile. We then checked from Sentinel-2 optical images for the conditions in each site where Emperor penguin guano can be detected (Fretwell and Trathan 2020).

## Methods

Geographic position of emperor penguin colonies from West Antarctica were downloaded from the Mapping Applicating for Penguin Populations and Projected Dynamics (MAPPPD ; Humphries et al. 2017; Lynch et al. 2022). A 50 km buffer around each of the 10 colonies (the minimum rounded distance between any of the colonies was ∼95km, therefore a 50km radius minimized buffer zone overlap) identified in the regions of the Amundsen and Bellingshausen Seas (between 70°W and 130°W ; Fig. 1), was used to download daily optimum interpolation of sea ice cover (SIC) data from ERDDAP server (Huang et al. 2023) between January 1, 1985 and June 20, 2023. Using the approach described in (Mora-Soto et al. 2022; Schlegel and Smit 2023), a past average SIC was calculated for each day of the year, using data from January 1, 1985 to December 31, 2022. Anomalies between daily observations and the 38-year SIC average were calculated between January 1, 2022 and June 20, 2023, and periods when SIC anomalies exceeded a value below the 25th percentile of the past average SIC for more than three consecutive days were identified as periods of extreme low sea ice cover. Events were classified as moderate (1 time below threshold) and strong (two times below threshold). Code for the analysis can be found at (Krüger 2023).

**Fig. 1.**
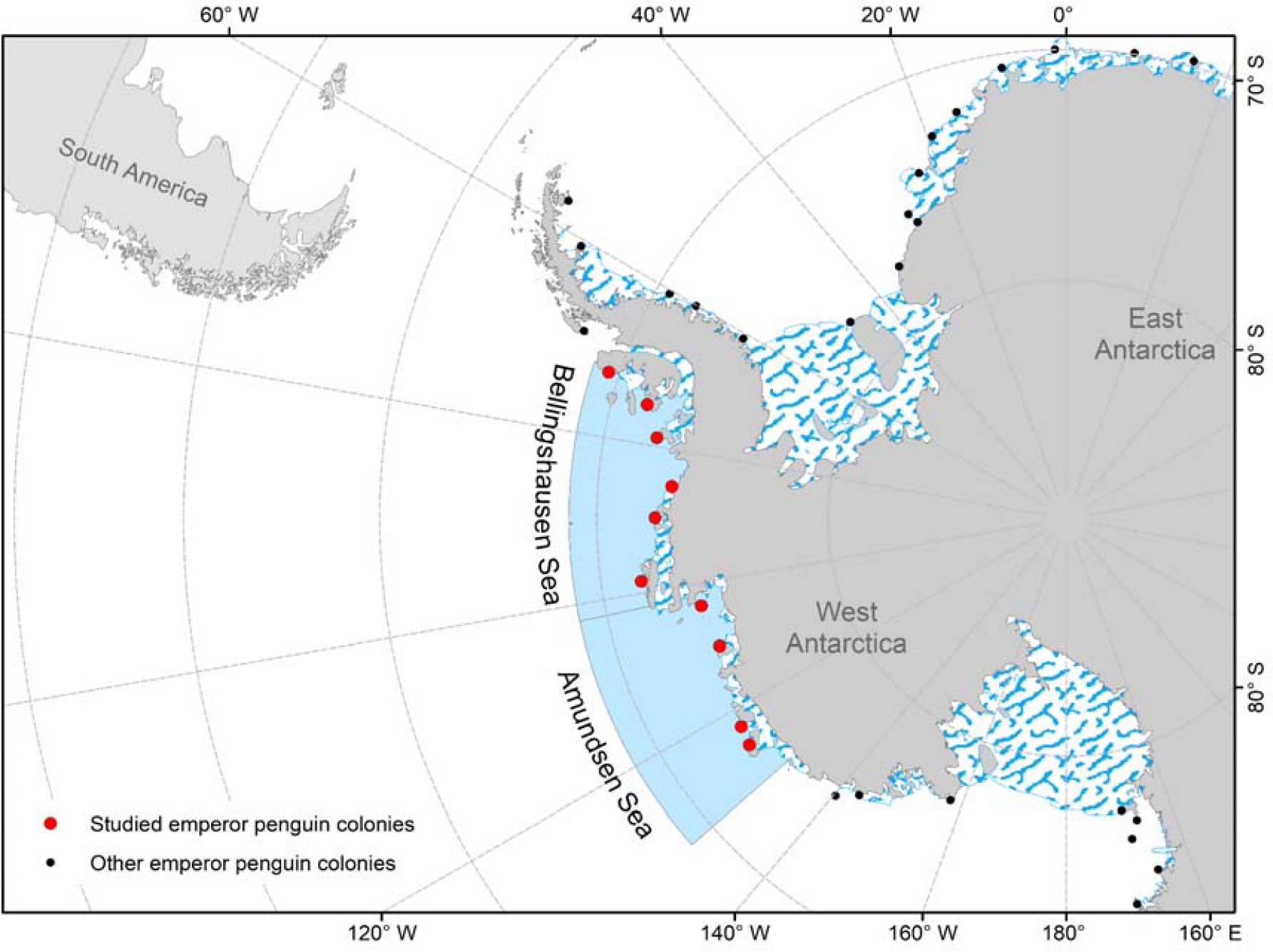
Location of studied emperor penguin colonies in the Bellingshausen and Amundsen Seas, West Antarctica. The geographic position of emperor penguin colonies was downloaded from the Mapping Application for Penguin Populations and Projected Dynamics (Humphries et al. 2017, Lynch et al. 2023). Limits of the Bellingshausen and Amundsen Seas adapted from (Larter et al. 2014; Christie et al. 2023).

Using the methods proposed in (Fretwell and Trathan 2020), freely available Sentinel-2 optical images (Sentinel-2 2023) were inspected using a cloud cover threshold of 50% and a custom rendering of bands that highlights penguin guano on the surface of the sea ice ([B08*0.8,B04*0.8,B03*0.8] Fretwell and Trathan 2020), between October and March of 2021/22 and 2022/23 breeding seasons. L1C or L2A (29 m x 29 m pixel size) images were downloaded. Guano stains were identified using an unsupervised classification algorithm in ArcMap.

## Results

The three colonies to the east of 80°W, in the Bellingshausen Sea(Rothschild Island, Verdi Inlet, Smyley Island), experienced short events (less than 15 days) of low sea ice cover between 2022 March and August, followed by longer events in October and November 2022 that persisted for over 30 days, and again in March to May 2023 (Fig. 2). The three colonies to the west of 80°W, in the Bellingshausen Sea (Bryan Coast, Pfrogner Point and Noville Peninsula), experienced strong and long events of low sea ice between November 2002 and January 2023 that persisted until middle May/early June (Fig. 2). For three colonies in the Amundsen Sea (west of 100°W; Brownson Island, Cape Gates, and Thurston Glacier), low sea ice events occurred in November and December 2022, January, February and March 2023, with two short low sea ice events at Brownson Island in June and July 2022 (Fig. 2). The colony at Bear Peninsula had a single short, low-intensity event in February 2023(Fig. 2).

**Fig. 2.**
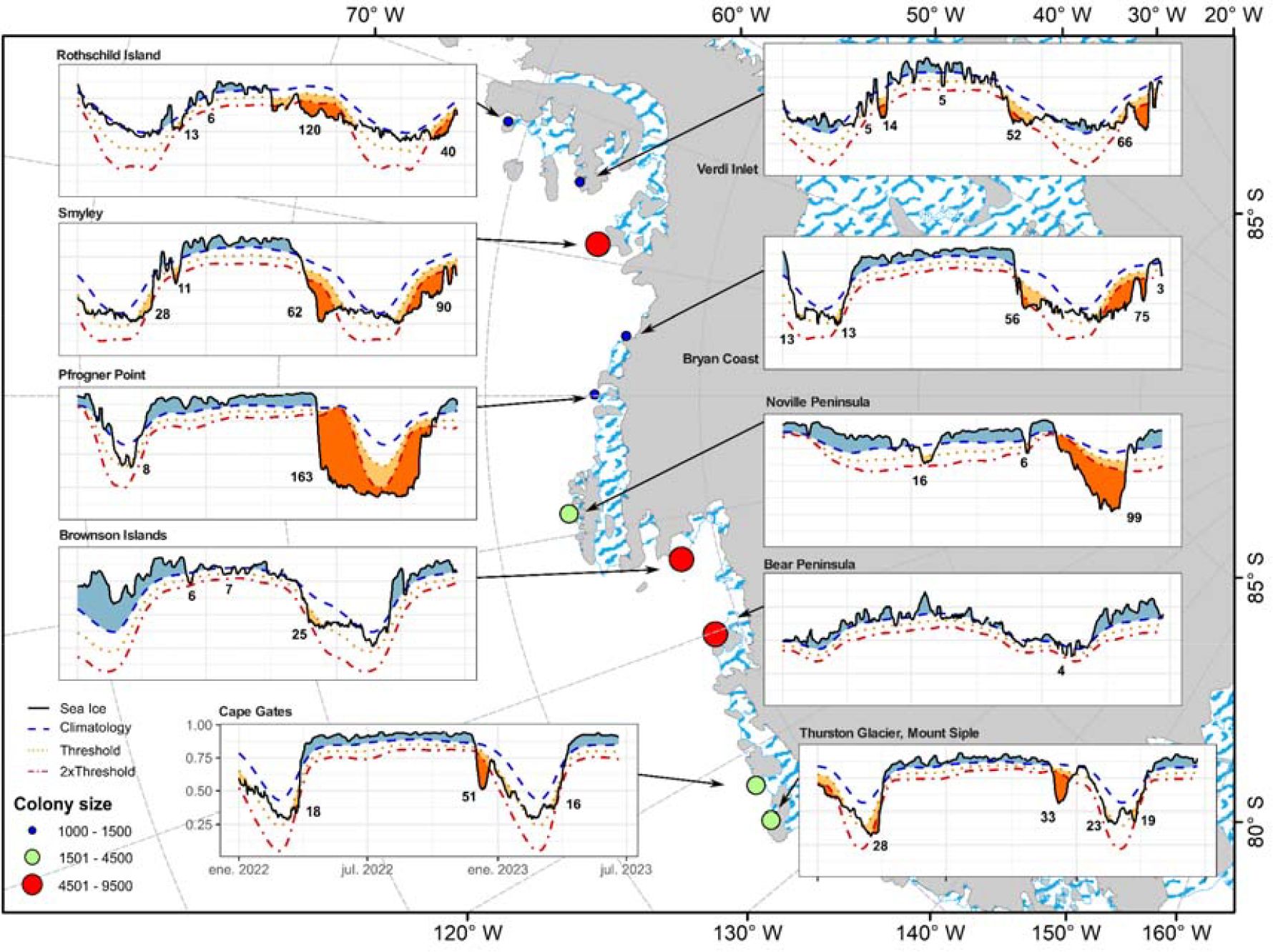
Emperor penguin colony sites (circles) in the Bellingshausen and Amundsen Sea region with estimated number of penguins (colony size) from the Mapping Application for Penguin Populations and Projected Dynamics (Humphries et al. 2017, Lynch et al. 2023). Daily sea ice cover within a 50 km radius around each colony, between January 1, 2022 and June 20, 2023 (solid black line in panels), is plotted against the 38-year SIC average (dashed blue line in panels),the 25th percentile critical threshold (dotted orange line in panels) and two times the threshold (dot-dash red line in panels) to identify low sea ice anomalies compared to historical values. Blue areas show periods when average sea ice cover was above the 38-year SIC average; orange areas show periods when average sea ice cover was below the critical (25th percentile) thresholds; and red areas show periods when negative anomalies were below 2x the critical thresholds. Numbers indicate days when sea ice cover remained below the critical threshold.

These episodes of low sea ice cover have affected the availability of sea-ice habitat at colony site upon the return of penguins to the breeding colonies in 2023 in 9 sites (Fig 3a-j) and during chick rearing of 2022 in 6 of the 10 sites (Fig3 a,b,c,d,e,g). In Brownson Islands (Fig 3g), it was possible to see the sea ice receding substantially around the crèches between November 20 and 30, 2022, which were then “trapped” in a small bay when there was no more sea ice on December 20, 2022. In Bear Peninsula (Fig 3h) the colony that was previously located over fast ice in 2020 moved into the sea shelf in 2022, and that shelf completely collapsed by March 2023. Thurston Glacier colony (Fig 3j) was the only site to have stable ice throughout 2022 and 2023.

**Fig. 3.**
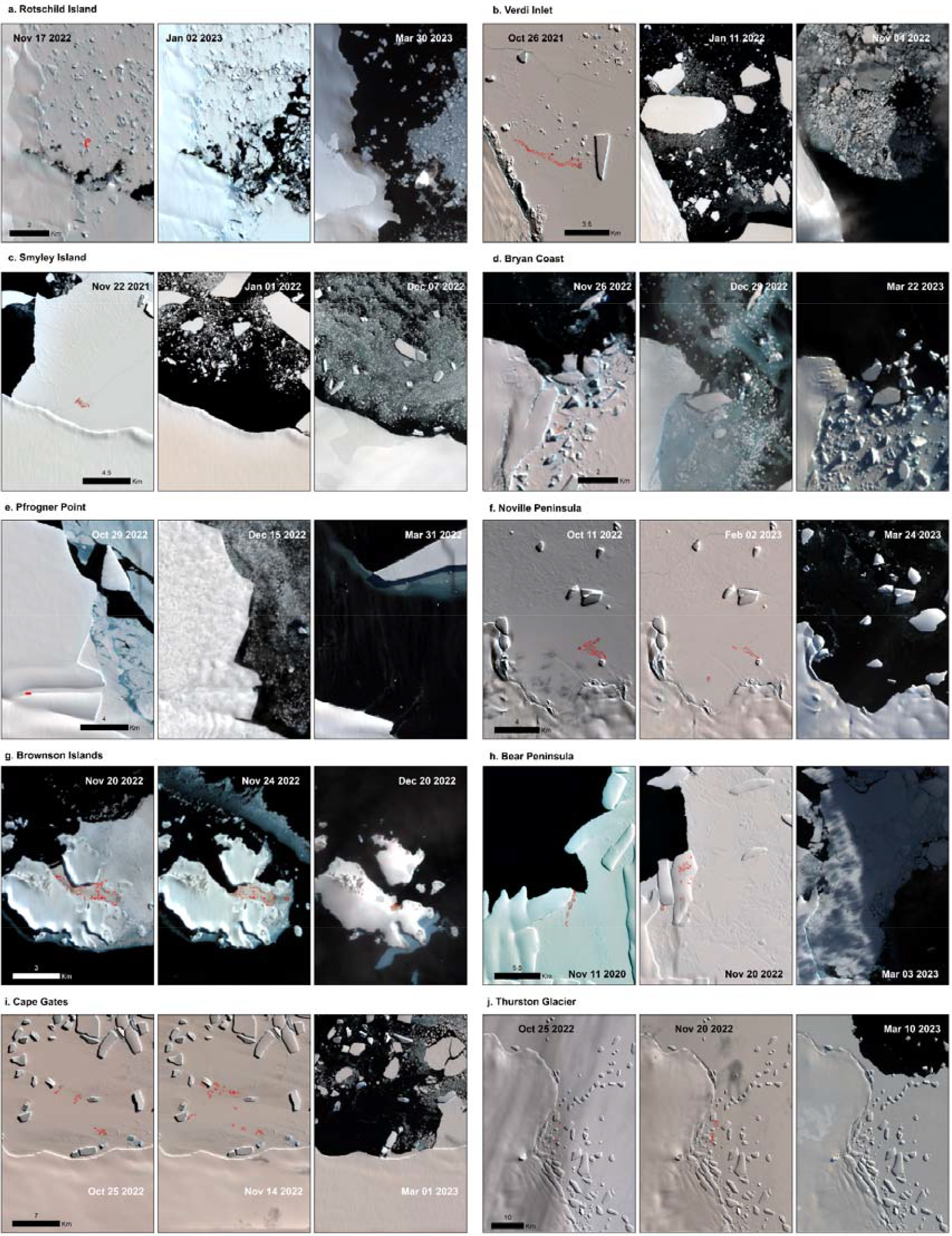
Sentinel-2 (2023) optical images at eight emperor penguin colonies from West Antarctica. Images used in the analyses consisted of recent images with less than 50% of cloud coverage in late crèche stage (October/November), during chick fledging period (December/January), and at the start of the breeding season (March/April), during episodes of low sea ice cover in 2022 and 2023. Red polygons (colony contours) were identified by unsupervised classification as penguin guano. The locations of the colonies presented here can be seen in Fig. 1.

## Discussion

Recent levels of sea ice cover in West Antarctica during key periods of emperor penguin breeding activity (the late chick-rearing period and the start of the breeding season) over the past two years have been lower than the 25th percentile of the last 38 years. For colonies in which satellite optical images were available for these periods, sea ice within the 50 km radius turned out to be critical at the breeding site.

While it is not possible to ascertain whether there has been breeding failure due to early sea ice retreats, sea ice losses prior to mid-December can result in complete breeding failure (Fretwell and Trathan 2019). There are examples of colonies that disappeared or reduced substantially in numbers after years of unsuitable breeding conditions, such as in Marguerite Bay, in the West Antarctic Peninsula (Trathan et al. 2011), and Halley Bay, east of the Weddell Sea (Fretwell and Trathan 2019). In addition, since the satellite imagery investigations began, several colonies have been known to “blink”, meaning that they disappear in some years, only to reappear in others (see Fig. 1 in Jenouvrier et al. 2021). It has been suggested that this could be a result of birds’ decision to move elsewhere after multiple consecutive failures or due to consecutive absence of optimal sea ice prior to arrival at the colony.

In the 2022/2023 season, sea ice extent was substantially low at most of the studied colonies in the Bellingshausen and Amundsen Seas in mid- to late-March (Fretwell et al. 2023, this study), when adults start returning to the colony to breed (Stonenhouse 1953; Prévost 1958). Adults may decide to skip breeding after arriving in suboptimal sea ice conditions (Kooyman and Ponganis 2017; Garnier et al. 2023), and there are examples of colonies that relocated to nearby ice shelves, as occurred for the colonies at Pfrogner Point (Trathan et al. 2020), Bear Peninsula (this study), Jason Peninsula and Atka Bay in the Weddell Sea (Fretwell et al. 2014; Zitterbart et al. 2014) or on the mainland for the Pointe Géologie colony (during the last 2022 and 2023 breeding seasons, Le Bohec unpublished observations). However, successive years of poor sea ice conditions can lead to site abandonment, as is assumed to have happened to colonies in Marguerite Bay, on the western Antarctic Peninsula (Trathan et al. 2011) and Halley Bay, on the eastern Weddell Sea (Fretwell and Trathan 2019).

Enhanced monitoring of emperor penguin colonies in the West Antarctic is paramount to assess, over the coming seasons, whether poor sea ice conditions persist and whether breeding colonies continue to occupy the areas. Satellite data with better resolution could be combined with field research (see Winterl et al. 2023) to assess the mortality associated with these events and its consequences on population trends.

### Conservation considerations

Recent studies have identified 29 Important Bird and Biodiversity Areas (IBAs) for emperor penguins distributed along the Antarctic coast, of which around 6 are located in the Bellingshausen and Amundsen Sea regions (Handley et al. 2021). Four of them (Smyley Island, Noville Peninsula, Brownson Island and Cape Gates) experienced sea ice loss during chick rearing or when adults returned to start a new breeding season, which likely represents a substantial proportion of the region’s breeding population.

Persistence of these colonies under expected changes in sea ice (Eayrs et al. 2021; Casagrande et al. 2023), depends on seabirds’ ability to disperse to new breeding sites. Garnier et al. (2023) suggested that emigration rates of emperor penguins are low and that their dispersal distance is short when compared to their post- and pre-breeding movements. They also suggested that the establishment in a new colony is random and that, when moving to a new colony, penguins may have limited knowledge of the sea ice conditions at the new site. Given the levels of similarity and spatial scales found in this study, even if emigration occurs in the Bellingshausen and Amundsen Sea regions, it will most likely be to sites with similar sea ice conditions. At-sea tracking data (non-existent for these colonies, see Houstin et al. 2022) combined with molecular techniques (i.e. Cristofari et al. 2016; Garnier et al. 2023) will be crucial for understanding the process of breeding site selection by new breeders and dispersal of adults under changing conditions.

Risks of local extinction for colonies in the Bellingshausen and Amundsen Sea region have been predicted to be high in the future, even with a forecasted 1.5°C increase in global temperature (Jenouvrier et al. 2021). The next few years will be of crucial importance in verifying whether emperor penguins will have the ability to persist in the region.

## Acknowledgments

This work was supported by the Instituto Antártico Chileno Programa Áreas Marinas Protegidas (AMP 24 03 052), ANID – Programa Iniciativa Milenio – ICN2021_044 (CGR) and ICN2021_002 (BASE).

